# Effect of interictal epileptiform discharges on EEG-based functional connectivity networks

**DOI:** 10.1101/784298

**Authors:** Derek K. Hu, Daniel W. Shrey, Beth A. Lopour

## Abstract

**Objective:** Functional connectivity networks (FCNs) based on interictal electroencephalography (EEG) can identify pathological brain networks associated with epilepsy. FCNs are altered by interictal epileptiform discharges (IEDs), but it is unknown whether this is due to the morphology of the IED or the underlying pathological activity. Therefore, we characterized the impact of IEDs on the FCN through simulations and EEG analysis.

**Methods:** We introduced simulated IEDs to sleep EEG recordings of eight healthy controls and analyzed the effect of IED amplitude and rate on the FCN. We then generated FCNs based on epochs with and without IEDs and compared them to the analogous FCNs from eight subjects with infantile spasms (IS), based on 1,340 visually marked IEDs. Differences in network structure and strength were assessed.

**Results:** IEDs in IS subjects caused increased connectivity strength but no change in network structure. In controls, simulated IEDs with physiological amplitudes and rates did not alter network strength or structure.

**Conclusions:** Increases in connectivity strength in IS subjects are not artifacts caused by the interictal spike waveform and may be related to the underlying pathophysiology of IS.

**Significance:** Dynamic changes in EEG-based FCNs during IEDs may be valuable for identification of pathological networks associated with epilepsy.

**Highlights:**

- Infantile spasms subjects exhibit broadly increased connectivity strength during interictal spikes
- Functional connectivity network structure is unaltered by interictal spikes in infantile spasms
- Simulated spikes in healthy control EEG did not alter network strength or structure

## 1. Introduction

Functional connectivity is a brain mapping technique based on the statistical interdependencies of spatially-distinct time-varying neural signals. Functional connectivity networks (FCNs) can provide valuable information about cortical network organization in both healthy subjects and those with epilepsy (Van Den Heuvel and Pol 2011; Kramer and Cash 2012; Van Diessen et al. 2013). Of the wide variety of imaging modalities used for functional connectivity, scalp electroencephalogram (EEG) is desirable due to its accessibility, low cost, standardized clinical application, noninvasive nature, and high temporal resolution. EEG-based FCNs have been used to characterize pathological networks associated with temporal lobe epilepsy (Quraan et al. 2013), benign epilepsy with centrotemporal spikes (Clemens et al. 2016; Mahmoudzadeh et al. 2016), and generalized pharmacoresistant epilepsies (Horstmann et al. 2010). When these FCNs are based on cross-correlation or coherence techniques using at least 100 seconds of EEG data, the networks exhibit stability over time, making them suitable for assessing an underlying disease state (Chu-Shore et al. 2012). For example, in infantile spasms (IS) subjects, strong, stable FCNs were found to underlie the chaotic EEG waveforms associated with hypsarrhythmia (Shrey et al. 2018).

In EEG-based FCNs, the inherent non-stationarity of the signal remains a challenge for analysis. For example, interictal epileptiform discharges (IEDs) are transient electrographic events that occur intermittently between seizures and are frequently recorded by EEG (de Curtis et al. 2012). In recent studies, IEDs have been shown to be correlated with a subject’s FCN, suggesting that the alteration of the baseline functional network may reflect pathological activity (Ponten et al. 2009; Horstmann et al. 2010; Adebimpe et al. 2015; Coito et al. 2016). While this prior work demonstrated that changes in FCNs occur during an IED, it is unknown whether these connectivity changes are driven purely by the morphology of the spike-wave complex (and its effect on the calculation of connectivity) or by the pathological network activity underlying the IED’s generation.

The goal of this study was to understand the effects of IEDs on FCNs from both methodological and physiological perspectives. Methodologically, we tested whether the presence of focal, high amplitude spike-wave complexes could cause spurious functional connectivity measurements. This was done by adding simulated focal IEDs at varying rates and amplitudes to the sleep EEG of control subjects and measuring the associated changes in the FCN. Once we understood the methodological effects of the IED waveform, we compared these results to the physiological changes in the FCNs derived from the sleep EEG of IS subjects exhibiting focal IEDs. IS subjects were chosen for this study due to the high epileptiform discharge amplitudes associated with this disease (Frost Jr et al. 2011). Based on prior EEG-based FCN studies, we hypothesized that the occurrence of focal IEDs would be associated with a local increase in functional connectivity strength (Wilke et al. 2011; Clemens et al. 2016) and the activation of a unique IED FCN (Ponten et al. 2009; Horstmann et al. 2010).

## 2. Methods

### 2.1 Subject information

Approval for this study was obtained from the Institutional Review Board of the Children’s Hospital of Orange County (CHOC), with the requirement for informed consent waived. We retrospectively identified eight infants (7F, 1M, aged 10.3±6.4 months) who were diagnosed with new-onset epileptic spasms and underwent scalp EEG recording prior to treatment. We also retrospectively identified eight control subjects (5F, 3M, aged 10.5±6.8 months) who (1) had no known neurological disorders, (2) underwent routine EEG for clinical evaluation, and (3) had EEGs that were interpreted as normal by a board-certified pediatric epileptologist (DS). All subjects had EEG recordings performed at CHOC.

### 2.2 EEG acquisition and preprocessing

All EEG data was recorded by the Nihon Kohden EEG acquisition system, with nineteen scalp electrodes placed according to the international 10-20 system, at a sampling rate of 200 Hz. For each subject, one interictal segment of sleep EEG lasting at least fifteen minutes was selected for analysis. Artifacts caused by muscle activity, movement, and poor electrode contact were marked by a board-certified epileptologist (DS). All electronic data were deidentified and analyzed using custom MATLAB (Mathworks) scripts. All EEG data were filtered with a third-order Butterworth filter with zero-phase shift digital filtering from 0.5-55 Hz and re-referenced to the common average. The data were then windowed into one-second epochs for connectivity analysis. Any epoch containing a marked artifact was discarded for all EEG channels after filtering. This data analysis procedure, including the subsequent calculation of functional connectivity, is summarized in Figure 1.

**Figure 1.**
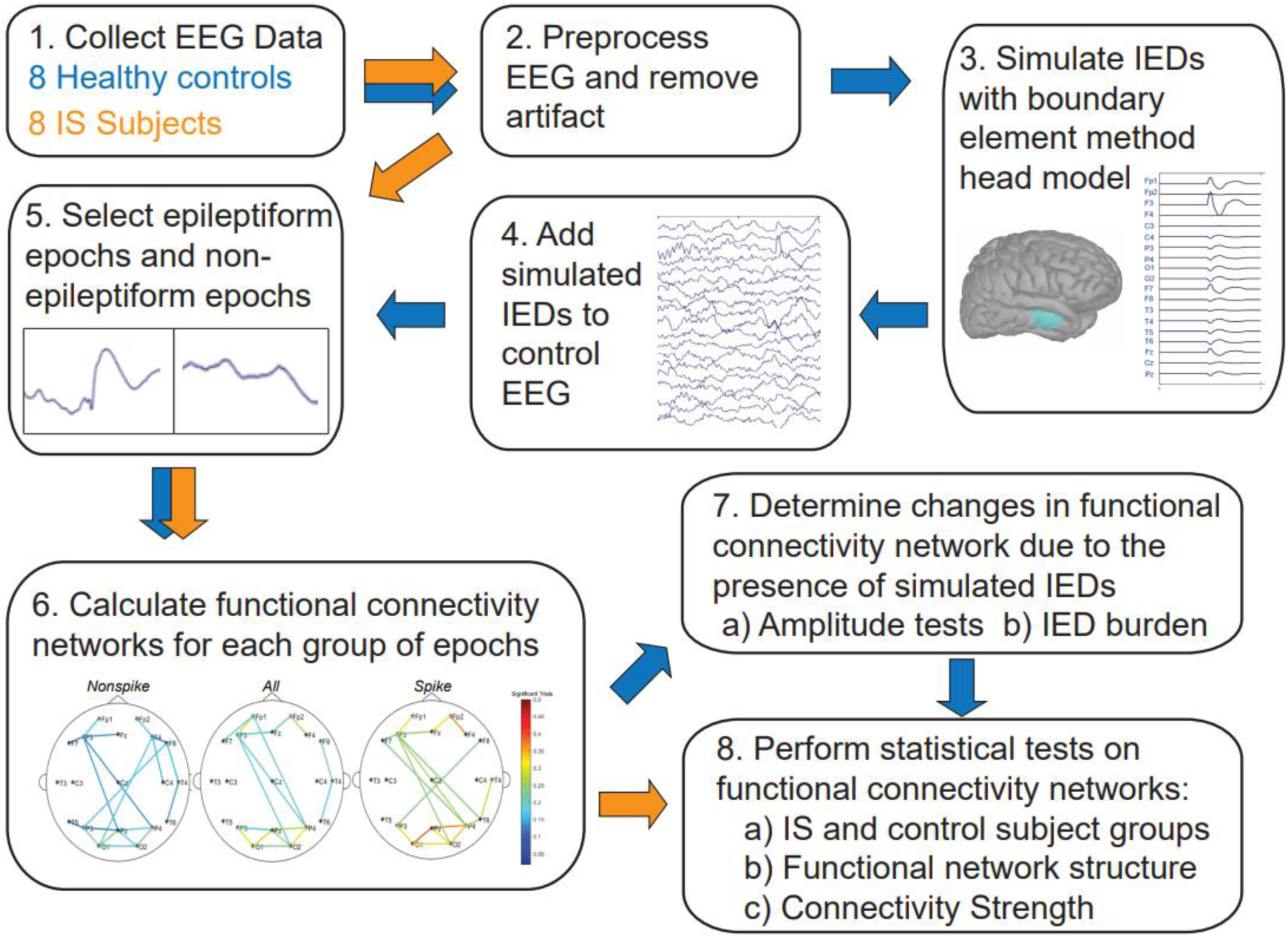
Summary of the functional connectivity analysis. Blue arrows indicate the data analysis procedure for the control subject EEG, while orange arrows indicate the procedure for analysis of EEG data from subjects with epilepsy.

### 2.3 EEG data segmentation for IS subjects

For each IS subject, the EEG data was segmented into three different groups: (1) all one-second epochs (ALL), (2) one-second epileptiform epochs (EE) containing an IED, and (3) one-second non-epileptiform epochs (NEE) containing no IEDs (Figure 2A). Focal IEDs were manually marked by a board-certified epileptologist (DS) based on the waveform morphology and the local field in adjacent electrodes. For each subject, the EEG channel containing the largest number of IEDs was selected for analysis. One epileptiform epoch was created for each IED within that channel, defined as the EEG data from all electrodes in a window of [−500, 500] milliseconds, centered on the spike. Each epoch was visually inspected to ensure that it contained only a single spike. The spike amplitude was defined as the EEG range within a [−50, 50] millisecond window centered around the spike, while background amplitudes were defined as the mean range of four 100-millisecond windows prior to the spike in each EE ([−500, −400], [−400, −300], [−300, −200], and [−200, −100] milliseconds). Non-epileptiform epochs were visually marked as one-second of EEG containing no epileptiform discharges in any channel.

**Figure 2.**
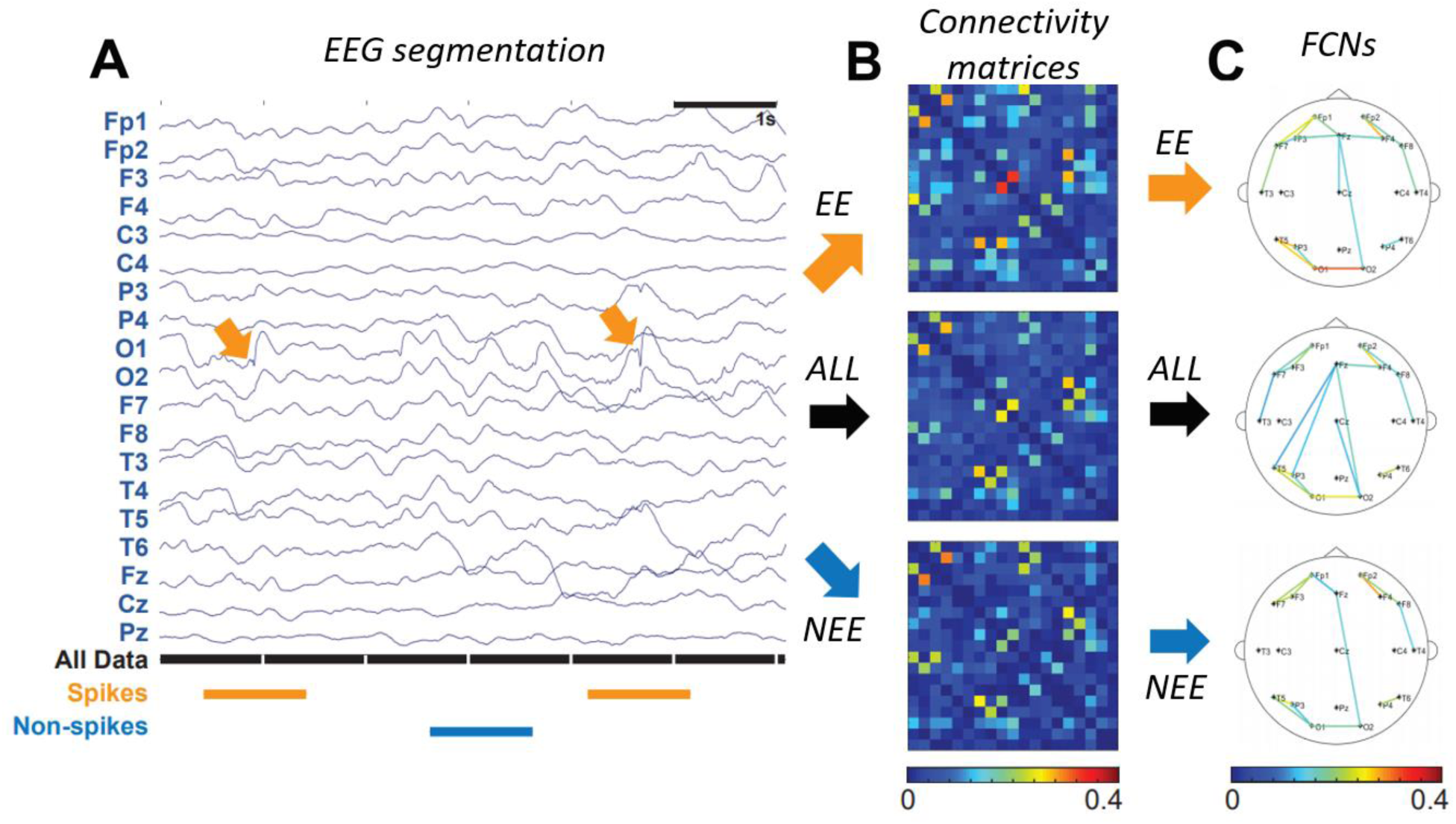
**(A)** Sample of EEG data segmentation. Orange segments are 1s epochs containing an IED (termed EE), black segments are all 1s epochs (ALL), and blue segments are 1s epochs with non-spike activity (NEE). **(B)** Connectivity matrices for spikes (orange arrow), all data (black arrow), and non-spikes (blue arrow), represented as a percentage of significant connections. **(C)** FCNs for EE (orange arrow), ALL (black arrow), and NEE (blue arrow). For clarity, only the strongest 10% of all connections are shown.

### 2.4 Simulation of IEDs in control subject EEGs

To assess the effect of IED waveforms on the FCN calculation, we simulated IEDs and added them to control sleep EEG recordings. Simulated IEDs were generated within a realistic head model using the Brainstorm software (Tadel et al. 2011). The realistic head model was based on the magnetic resonance imaging template brain volume in the ICBM 152 atlas (Fonov et al. 2011). The atlas was used with boundary element methods (BEM) to generate a three-layered geometric head model, consisting of the scalp, inner skull, and outer skull with a conductivity ratio of 1:0.0125:1, respectively.

IED simulations were based on the equivalent current dipole method, where it is assumed that the spike potential is generated from a primary dipole source (Koles 1998; Grova et al. 2006). Here, we chose a single patch of activated cortex located underneath the F3 electrode, with the corresponding dipole oriented normal to the selected region. The region of activated cortex was 7-10 cm^2^, concordant with the typical surface area of cortex necessary to produce epileptiform discharges that are detectable on scalp EEG (Grova et al. 2006; Tao et al. 2007). We then modelled the time course of the spike-wave complex as a combination of three half-period sine waves within a one-second window. The simulated spike was represented by a positive half-period sine wave 60 milliseconds long, followed by a slow wave consisting of sequential negative and positive half-period sine waves lasting 120 milliseconds and 200 milliseconds, respectively (Figure 3). The amplitude ratio for each half-wave was 5:4:2, which was chosen to approximate the average characteristics of IEDs marked in IS subjects. The IED field across all nineteen electrodes was computed using forward modelling with the OpenMEEG software, incorporating the generated simulated spike waveform and geometric head model.(Kybic et al. 2005; Gramfort et al. 2010). To simulate the sporadic occurrence of IEDs, the IED waveform was added to a randomly selected subset of nonoverlapping one-second EEG epochs, with the peak of the spike aligned to the center of each epoch.

**Figure 3.**
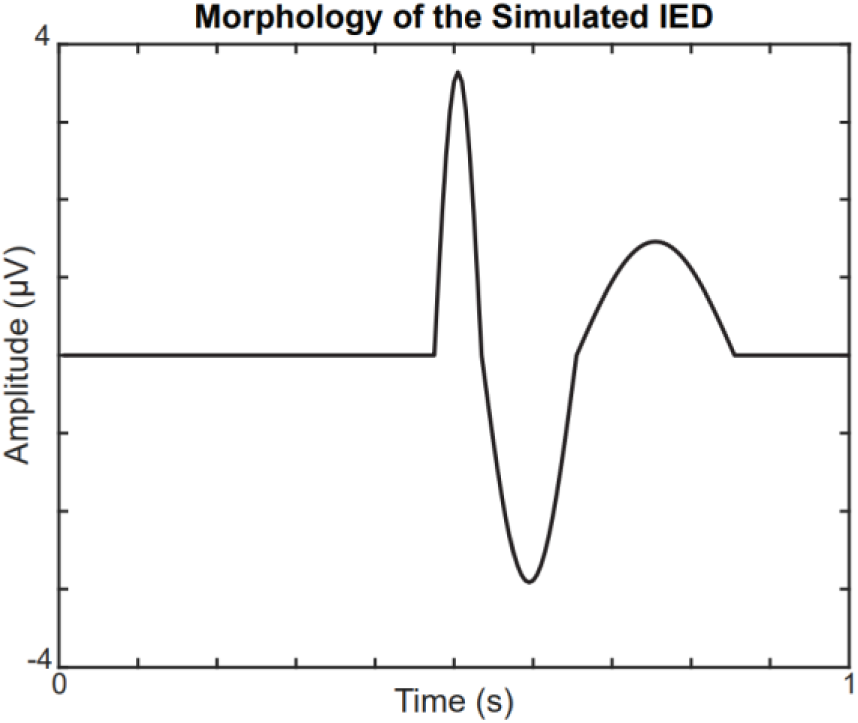
Morphology of the simulated IED at the focal electrode F3. The amplitude of the IED is scaled to match the spike:background ratio of IS subjects.

### 2.5 Calculation of the functional connectivity network

For each subject, FCNs were calculated using linear cross-correlation, as previously described in (Kramer et al. 2009; Chu-Shore et al. 2012; Shrey et al. 2018). This technique has been shown to provide accurate measurements for both real and simulated electrophysiological data (Jiruška et al. 2005) and generate stable FCNs when using at least 100 seconds of data (Chu-Shore et al. 2012). Prior to FCN analysis, the data in each one-second epoch was normalized in each channel to have zero mean and unit variance. To calculate the coupling strength between each electrode pair, we first calculated the maximal absolute value of the cross correlation with a maximum lead/lag time of 200 milliseconds. This maximum lead/lag was chosen based on typical times for neurophysiological processes and cross-cortical conduction times (Chu-Shore et al. 2012). Connectivity values were normalized based on the autocorrelation of the signal at the chosen lead/lag time. We then applied a Fisher z-score transformation, resulting in an adjacency matrix containing the z-score between each electrode pair for each one-second epoch.

The significance of the coupling for each one-second epoch was determined by comparing each electrode pair’s z-score value to a null distribution generated using permutation resampling. In each iteration of resampling, we computed the maximal absolute value of the cross-correlation (with a max lead/lag of 200 milliseconds) between random one-second epochs for each channel, after excluding epochs containing artifacts. This was repeated 1000 times to create a normal distribution of z-scores for each electrode pair under the null hypothesis that there was no temporal relationship between the two channels. The cross-correlation in each one-second epoch was considered statistically significant if the computed z-score value was higher than the 95^th^ percentile of the null distribution.

To prevent spurious connections due to volume conduction, any z-score with a maximal cross correlation at a zero-time lag was considered non-significant (Chu-Shore et al. 2012). The results for each one-second epoch were stored in a binary adjacency matrix, where a value of one represented a significant, non-volume conducted connection. We then averaged the binary adjacency matrices to produce a connectivity matrix where each element represented the percentage of significant connections between the electrode pairs over the duration of the recording (Figure 2B). For visualization, we created topographical network maps by applying a threshold to the connectivity matrix (Figure 2C).

For each IS subject, three different FCNs were constructed using the ALL, EE, and NEE epochs segmented from the EEG. In control subjects, we generated one FCN from the control EEG before adding simulated IEDs (CONTROL) and three FCNs from control EEG with simulated IEDs using different subsets of epochs: (1) All epochs after adding simulated IEDs (ALL), (2) only epochs containing simulated focal IEDs (EE), and (3) only epochs not containing simulated IEDs (NEE). The same null distribution was used for calculating the FCNs of the ALL, EE, and NEE groups. To measure the variance of our results, we performed 250 iterations of FCN generation for each control subject, with each iteration containing simulated IEDs in randomly selected epochs.

### 2.6 Statistical tests for network structure

To quantify the differences in network structure between two FCN’s, we used two different metrics: the relative graph edit distance (rGED) and 2D correlation. The rGED is a novel method based on the principles of the graph edit distance that measures the similarity between two binary graphs with the same number of vertices and edges (E) (Sanfeliu and Fu 1983). The rGED is calculated based on the minimum number of insertions (I) and deletions (D) required to transform one of the graphs into the other one: 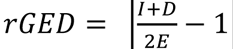. An rGED of zero indicates that there are no connections in common, while an rGED of one indicates that both networks have all connections in common. To calculate the rGED between two FCNs, we binarize each connectivity matrix by assigning the strongest ten percent of connections a “1” and all other connections a “0”.

The 2D correlation measures the similarities across the entire FCN rather than focusing on the strongest connections, as is done with the rGED. Using 2D correlation also obviates the need for thresholding, which can potentially bias the measurement. Both the rGED and 2D correlation tests were used to compare the ALL to EE, ALL to NEE, and EE to NEE networks in IS and control subjects. The 2D correlation was also used to compare the differences in network structure between IS and control FCNs. Specifically, we compared the ALL FCN of each IS subject to the ALL FCNs of all other IS subjects (n=28 comparisons), the CONTROL FCN of each control subject to the CONTROL FCNs of all other control subjects (n=28), and the ALL FCN of each IS subject to the CONTROL FCNs of all control subjects (n=64). Comparisons between these distributions show the uniformity of FCNs within each group, compared to comparisons across different groups.

### 2.7 Statistical tests for connectivity strength

To quantify the changes in connectivity strength during physiological IEDs in IS subjects, we performed statistical tests to compare an individual’s ALL, EE, and NEE FCNs. Based on previous studies reporting an increase in functional connectivity during IEDs (Siniatchkin et al. 2007; Wilke et al. 2011; de Curtis et al. 2012; Clemens et al. 2016), we expected the EE FCNs to have the highest connectivity strength, followed by the ALL FCNs, followed by the NEE FCNs. We tested these comparisons for each IS subject using three one-tailed Wilcoxon sign-rank tests, where each test compared different FCN pairs: 1) EE > ALL, 2) ALL>NEE, and 3) EE>NEE. In each test, we compared the paired distributions of all connections to the null hypothesis that the median strengths were not statistically different.

## 3. Results

### 3.1 Amplitude and burden of the simulated IEDs

We defined the simulated IED waveform’s amplitude and burden based on the spike to background amplitude ratio and the frequency of IEDs in IS subjects (see Section 2.3). Across all patients, our analysis included 1,340 visually marked IEDs and 5,360 measurements of background amplitude. The mean spike to background amplitude ratio across all eight IS subjects was 2.62 to 1, with a standard deviation of 0.38. To match the spike to background ratios seen in IS subjects, we scaled the simulated spike amplitude based on the average background amplitude across all controls (26.8±6.3 μV). In our simulations, we conservatively accounted for variance in the control subjects by using three standard deviations above the average of the control background amplitude (45.8 uV), multiplied by the spike:background ratio of 2.62, to generate 120 uV discharges. The spike burden for each subject was defined as the number of marked focal IEDs divided by the total EEG recording time (Table 1). The highest burden across all eight IS subjects was approximately 25%, so we conservatively used this maximum spike burden for all simulations, unless otherwise specified.

**Table I.**
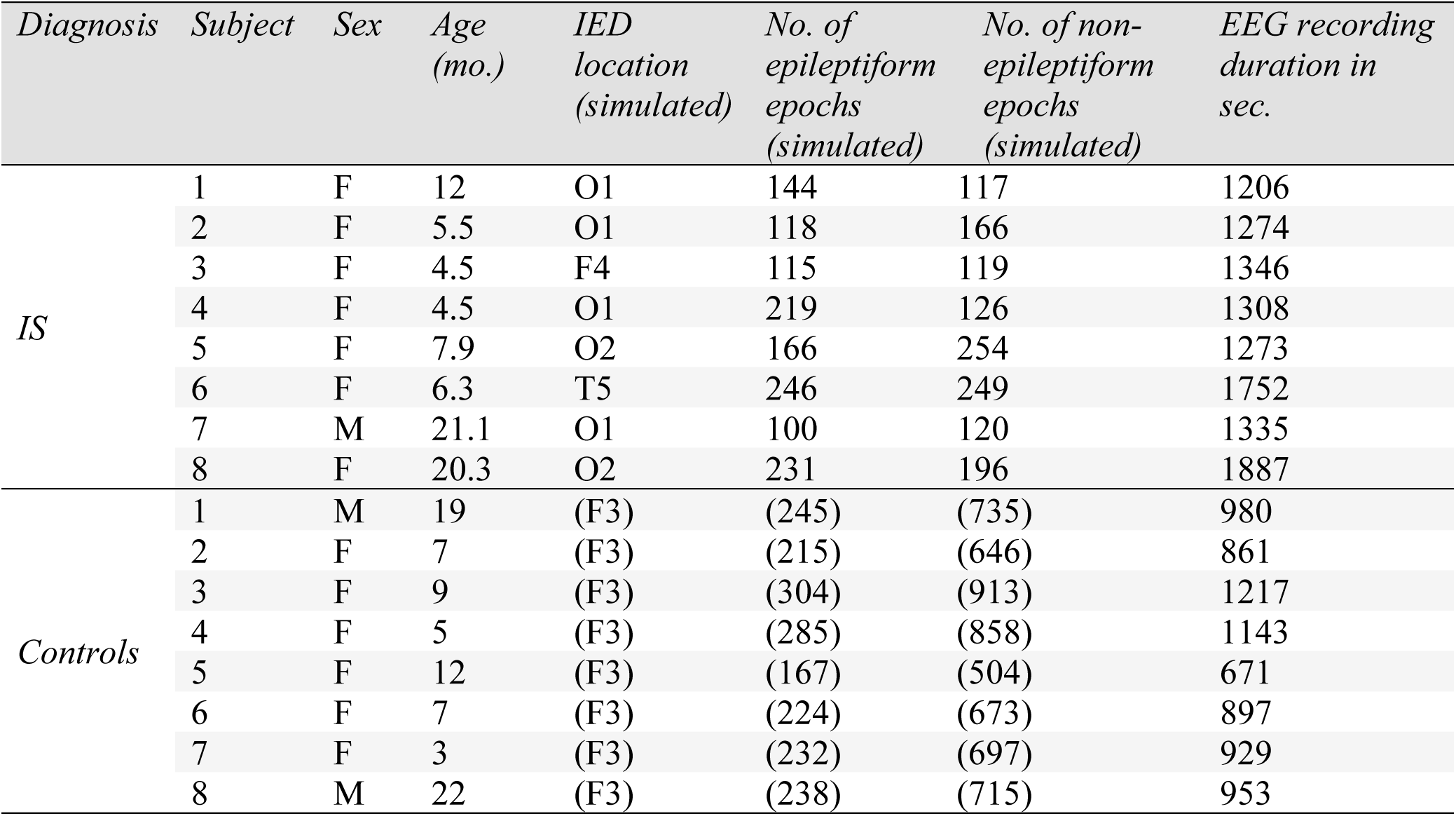
Patient demographics and clinical data

| *Diagnosis* | *Subject* | *Sex* | *Age*<br>*(mo.)* | *IED*<br>*location*<br>*(simulated)* | *No. of*<br>*epileptiform*<br>*epochs*<br>*(simulated)* | *No. of non-*<br>*epileptiform*<br>*epochs*<br>*(simulated)* | *EEG recording*<br>*duration in*<br>*sec.* |
| --- | --- | --- | --- | --- | --- | --- | --- |
| *IS* | 1 | F | 12 | O1 | 144 | 117 | 1206 |
|  | 2 | F | 5.5 | O1 | 118 | 166 | 1274 |
|  | 3 | F | 4.5 | F4 | 115 | 119 | 1346 |
|  | 4 | F | 4.5 | O1 | 219 | 126 | 1308 |
|  | 5 | F | 7.9 | O2 | 166 | 254 | 1273 |
|  | 6 | F | 6.3 | T5 | 246 | 249 | 1752 |
|  | 7 | M | 21.1 | O1 | 100 | 120 | 1335 |
|  | 8 | F | 20.3 | O2 | 231 | 196 | 1887 |
| *Controls* | 1 | M | 19 | (F3) | (245) | (735) | 980 |
|  | 2 | F | 7 | (F3) | (215) | (646) | 861 |
|  | 3 | F | 9 | (F3) | (304) | (913) | 1217 |
|  | 4 | F | 5 | (F3) | (285) | (858) | 1143 |
|  | 5 | F | 12 | (F3) | (167) | (504) | 671 |
|  | 6 | F | 7 | (F3) | (224) | (673) | 897 |
|  | 7 | F | 3 | (F3) | (232) | (697) | 929 |
|  | 8 | M | 22 | (F3) | (238) | (715) | 953 |

### 3.2 Excessive spike amplitudes change FCN structure and increase connectivity strength

To determine the effect of the simulated IED’s amplitude on a subject’s FCN, we added simulated IEDs with varying spike:background amplitude ratios to control subject EEG and compared the EE and ALL FCNs to the CONTROL FCN. The addition of simulated spikes had a small but significant effect on both the network structure and mean connectivity strength in the ALL FCN (Figures 4A, 5A, 5B) (Wilcoxon rank sum test, p<0.05 pre-specified threshold FDR, corrected for multiple comparisons using the Bonferroni correction; pFDR = 0.00714). In contrast, we saw large changes in both the network structure and mean connectivity strength in the EE FCN (Figures 4B, 5C, 5D) (Wilcoxon rank sum test, p<0.05 pre-specified threshold FDR, corrected for multiple comparisons using the Bonferroni correction; pFDR = 0.00714). Across all control subjects, we found a dramatic decrease in 2D correlation (indicating a change in network structure) and an increase in mean connection strength at spike:background ratios greater than 7, which is more than 2.5 times greater than the average spike:background ratio for IS subjects (Supplementary Figures 1-7).

**Figure 4.**
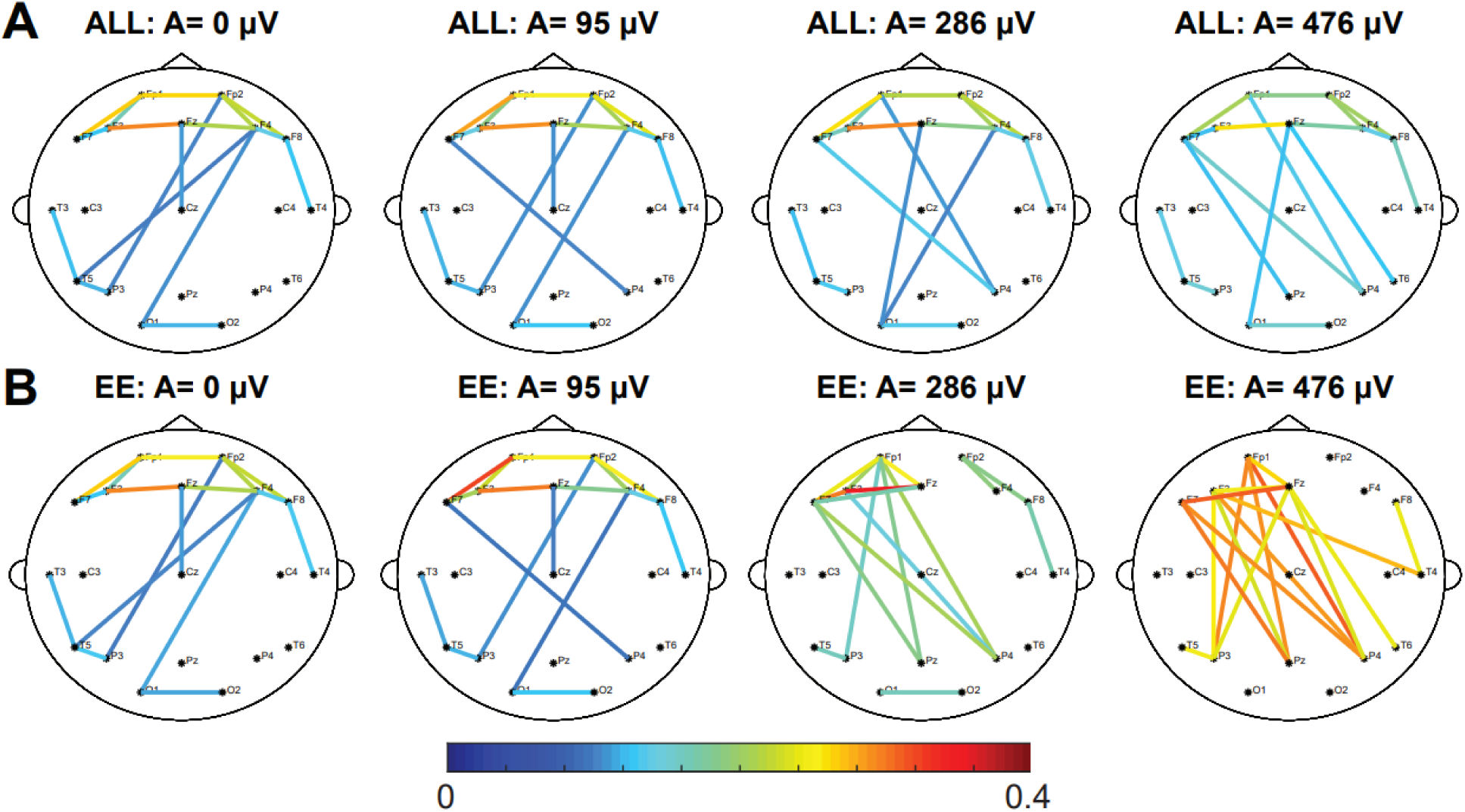
Increasing the amplitude of the IED has a small effect on the strength and structure of the ALL network and a large effect on the EE network. Representative example showing the effect of adding simulated focal IEDs at F3 with varying amplitudes on the (A) ALL FCN and the (B) EE FCN of control subject 6. The strongest 10% of connections are shown. Results for all other control subjects are shown in Supplementary Figures 1-7.

**Figure 5.**
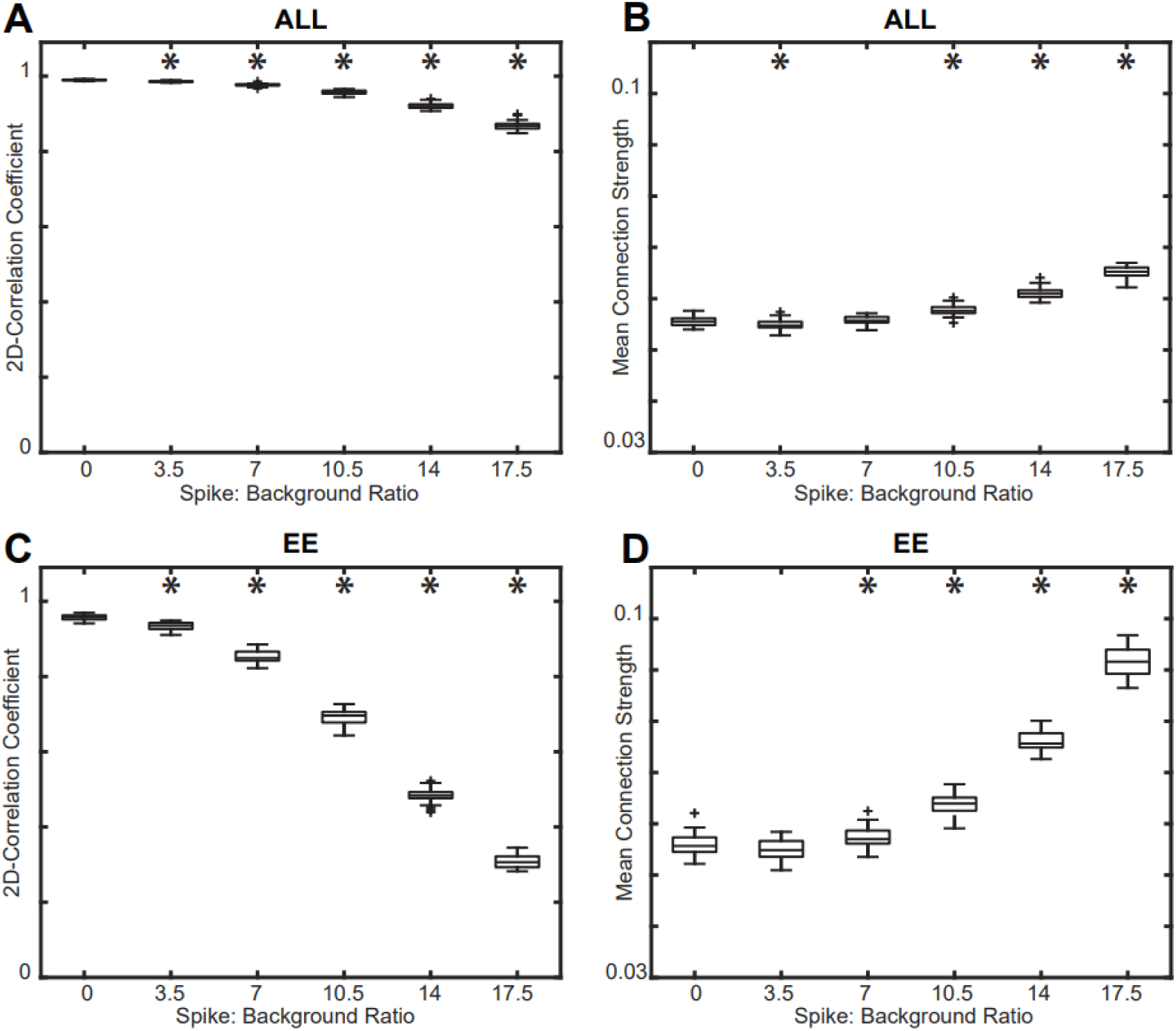
Effect of varying spike amplitude on the (A) 2D correlation and (B) mean connection strength of the ALL FCN compared to the CONTROL FCN. Effect of varying spike amplitude on the (C) 2D correlation and (D) mean connection strength of the EE FCN compared to the CONTROL FCN. The mean spike:background ratio for IS subjects was 2.62, and changes in network strength and structure occur well above this value. Results for all control subjects were similar (Supplementary Figures 1-7); a representative example from control subject 6 is shown. Significance tests compared each result to the base CONTROL FCN (spike burden = 0%).

### 3.3 Increasing the spike burden has little effect on connectivity strength

Next, we investigated changes in the FCN as a function of spike burden. We inserted varying numbers of IEDs into the control subject EEG; specifically, we added them to 0%, 20%, 40%, 60%, 80%, and 100% of all one-second epochs and then calculated the FCN (Figure 6). We found that increasing the number of IEDs slightly decreased the ALL FCN’s correlation to the CONTROL network from 0.990 to 0.949 (Figure 7A) and increased the mean connectivity strength from 0.0555 to 0.0575 (Figure 7B) (Wilcoxon rank sum, p<0.05 pre-specified threshold FDR, corrected for multiple comparisons using the Bonferroni correction; pFDR = 0.01). Although these differences in strength and structure were small, they were statistically significant due to the low variance across simulations (Figure 7). These results were consistent across all control subjects (Supplementary Figures 1-7).

**Figure 6.**
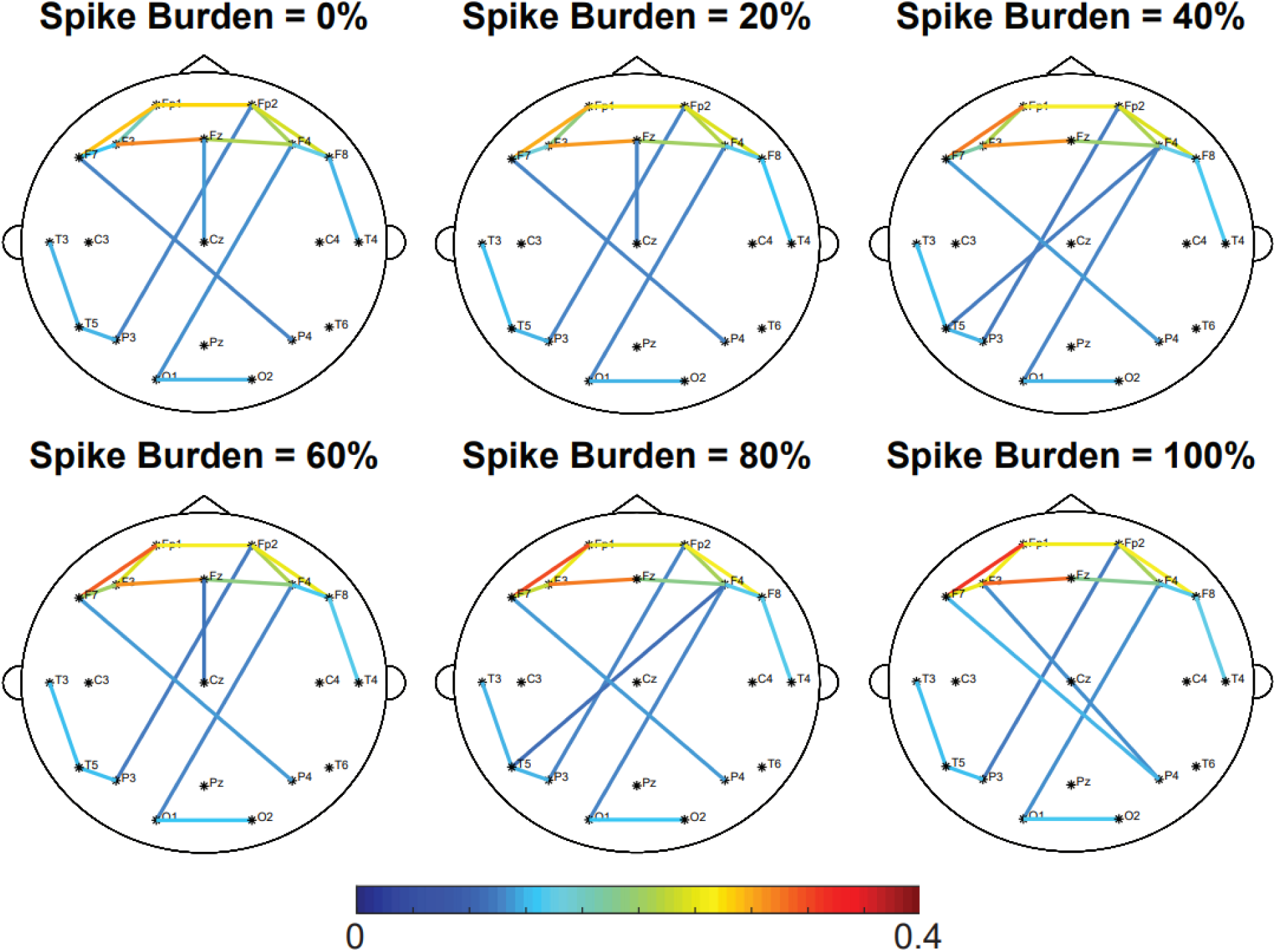
Increasing the spike burden does not affect the network strength and structure of the ALL network. Representative example of the effect of spike burden on the ALL FCN of control subject 6. Simulated focal IEDs are added at electrode F3. The strongest 10% of connections are shown. Results for all other control subjects are shown in Supplementary Figures 1-7.

**Figure 7.**
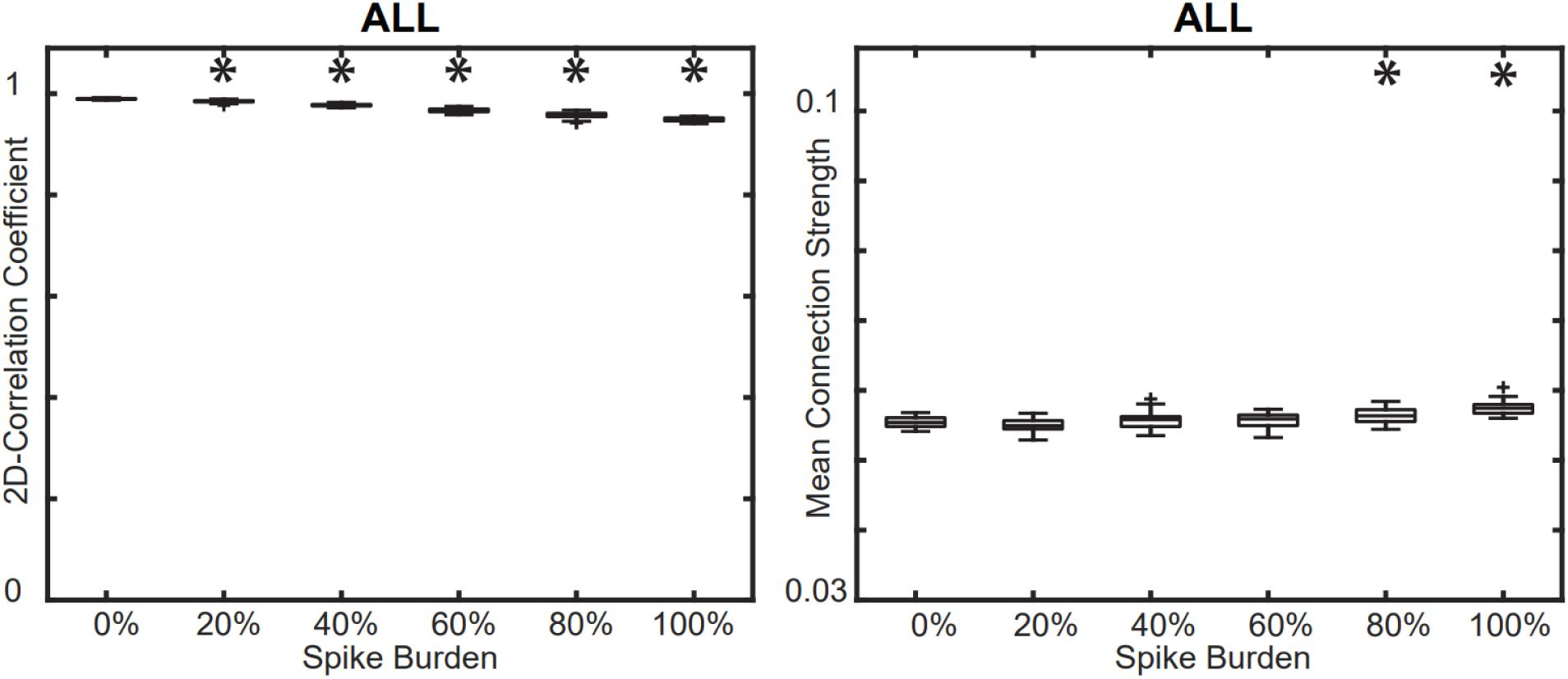
(A) Effect of spike burden on the 2D correlation between the ALL FCN and CONTROL FCN. (B) Effect of spike burden on the mean connection strength in the ALL FCN. Results for all control subjects were similar (Supplementary Figures 1-7); a representative example from control subject 6 is shown. Significance tests compared each result to the base CONTROL FCN (spike burden = 0%).

### 3.4 FCN structure for IS subjects is subject-specific

We then quantified the differences in the network structure of individual IS and control subjects. This was done by calculating the 2D correlation of the connectivity matrices between all pairs of subjects in the following within-group or across-group comparisons: (1) control-control, (2) IS-control, and (3) IS-IS. Figure 8 shows the correlation coefficients and statistical comparisons for these three categories (Wilcoxon rank sum, p<0.05 pre-specified threshold FDR, corrected for multiple comparisons using the Bonferroni correction; pFDR = 0.0167). Control-control correlation coefficients were significantly higher than those for the IS-control (p = 1.163e-7) and IS-IS comparisons (p = 1.064e-4). We found no significant differences in correlation between IS-IS and IS-control (p = 0.6196). These results suggest that the FCNs of control subjects are more stereotyped and exhibit less variability than the FCNs of IS subjects. Our findings also demonstrate that the FCNs of individual IS subjects are no more similar to other IS subjects than they are to control subjects.

**Figure 8.**
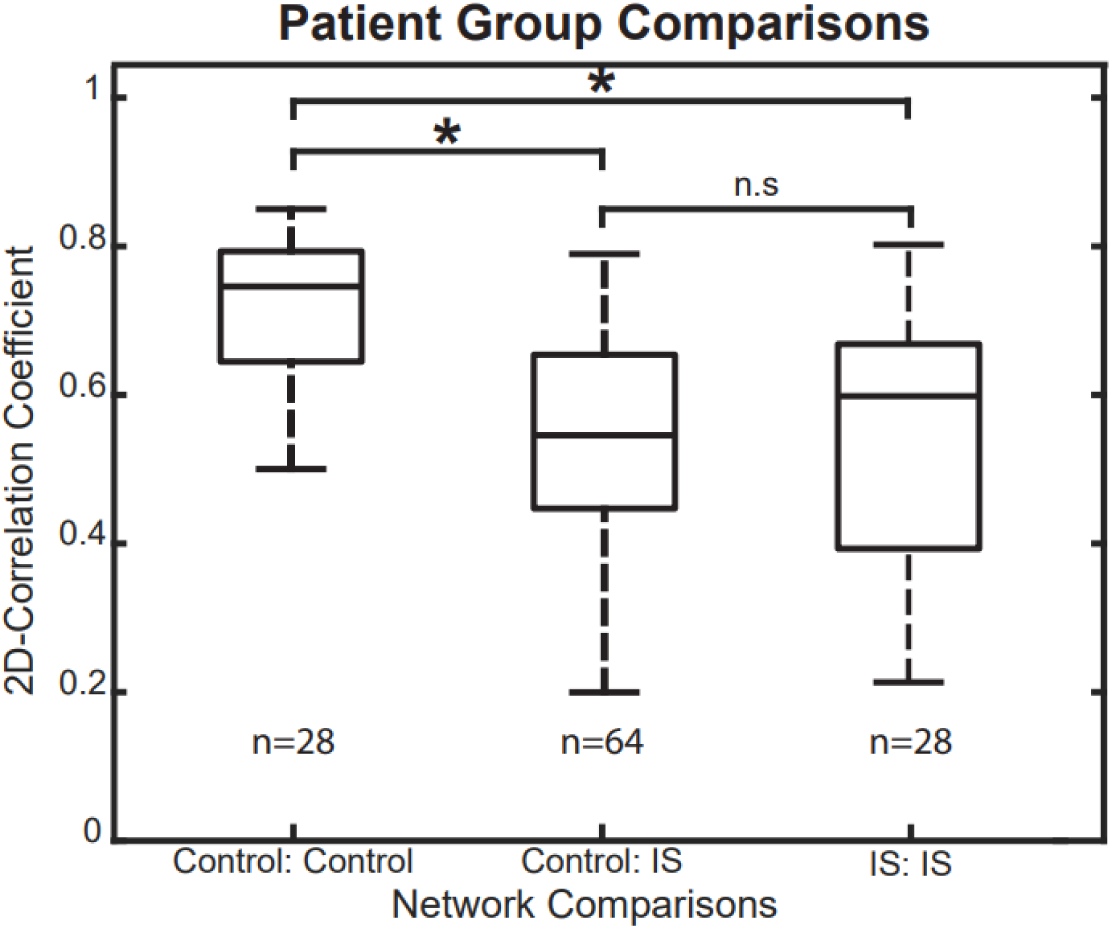
Rank-sum tests for 2D correlation coefficients across different subject groups. The correlation coefficients in control-control comparisons were significantly higher than in control-IS and IS-IS comparisons, indicating that control subjects have more consistent, stereotyped FCNs.

### 3.5 FCN structure does not change during interictal spikes

In both IS and control subjects, changes in network structure associated with the presence of IEDs were assessed using the rGED and 2D correlation, two complementary techniques that yielded similar results. We compared all three pairs of FCNs within an individual subject: (1) ALL compared to NEE, (2) ALL compared to EE, and (3) EE compared to NEE. To gain perspective on the degree of similarity between these networks, we compared these values to a null model consisting of network comparisons across different subjects within the same cohort. For example, to test the ALL vs. NEE FCNs of Subject 1, we calculated the rGED and 2D correlation of Subject 1’s ALL to Subject 1’s NEE FCN. These values were then compared to a null model consisting of rGED and 2D correlation values of Subject 1’s ALL vs. Subject 2’s NEE, Subject 1 ALL vs. Subject 3 NEE etc. plus the analogous correlations between Subject 1’s NEE to the ALL from all other subjects. If the intra-subject FCN correlations were higher than the inter-subject FCN correlations, this indicated that the presence of IEDs did not significantly alter the subject’s FCN.

For all eight IS subjects, the intra-subject rGED values for the ALL vs. NEE and ALL vs. EE network comparison tests were significantly higher than the inter-subject values (Figure 9A) (one sample Wilcoxon signed rank test, p<0.05 pre-specified threshold FDR, corrected for multiple comparisons using the Bonferroni correction; pFDR = .00625). In the EE vs. NEE network test, seven out of eight epilepsy subjects had significant rGED values. This demonstrates that the ALL, EE, and NEE FCNs are subject-specific and that the strongest connections are not significantly affected by the presence of IEDs. In control subjects with simulated IEDs, we found significantly higher intra-subject rGEDs compared to inter-subject rGEDs across the control subjects in all network comparisons across 200 different iterations of simulated IEDs, (Wilcoxon rank sum, n= 8, p<0.05 pre-specified threshold FDR, corrected for multiple comparisons using the Bonferroni correction; pFDR = 0.00625) (Figure 9B).

**Figure 9.**
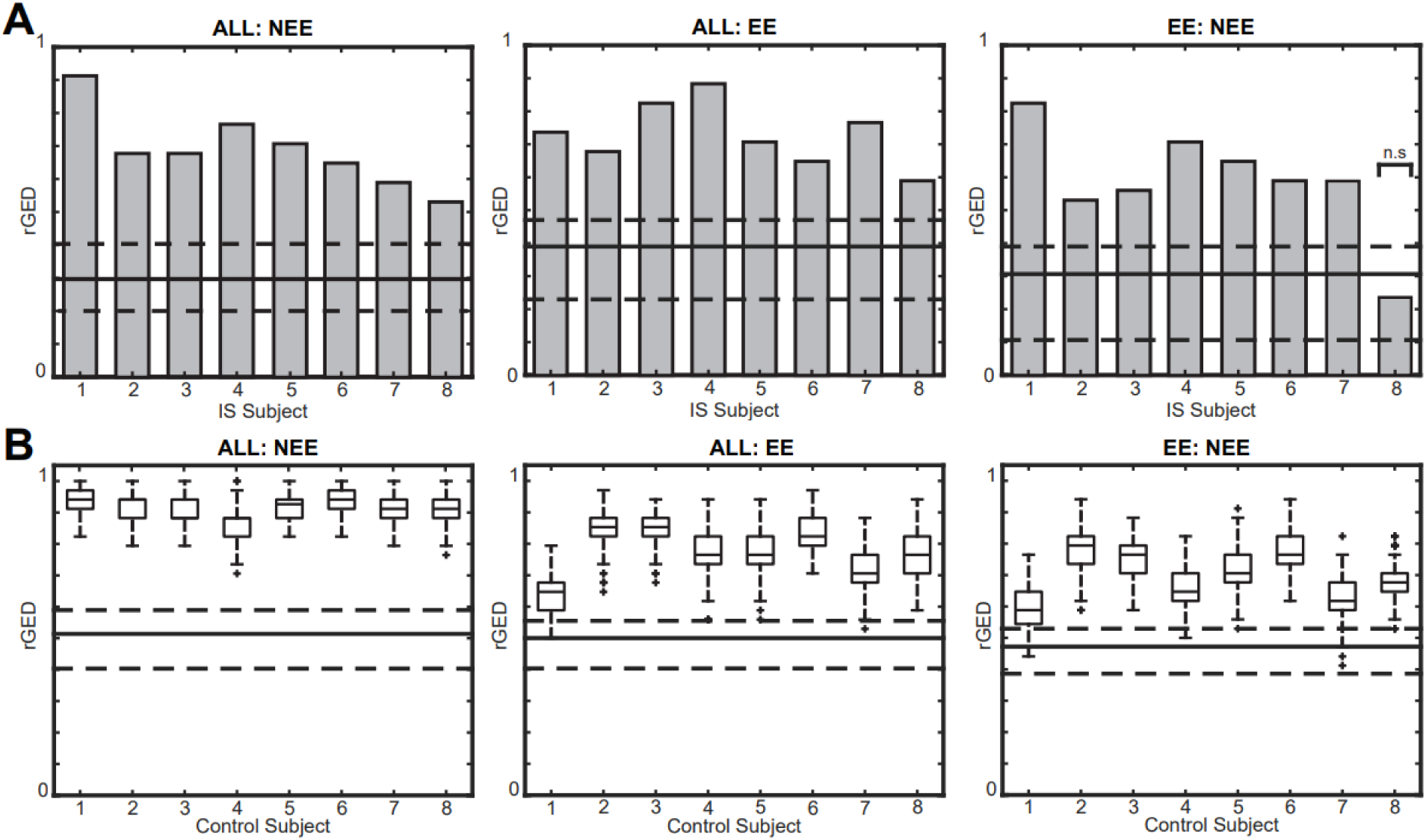
Relative graph edit distance (rGED) values for intra-subject network comparisons are significantly higher than inter-subject comparisons in (A) IS subjects and (B) control subjects with simulated IEDs. Gray bars represent intra-subject comparisons for IS subjects, and box plots represent intra-subject values from 200 simulations in controls. The solid lines represent the median of the inter-subject rGED values and the dashed lines represent the 25^th^ and 75^th^ percentiles. All tests are significant except EE:NEE in IS subject 8.

The analogous calculation using 2D correlation yielded similar results (Figure 10). These results were significant for all three network comparisons across all eight IS subjects (one sample Wilcoxon signed rank test, p<0.05 pre-specified threshold FDR, corrected for multiple comparisons using the Bonferroni correction; pFDR = .00625) and all eight control subjects (Wilcoxon rank sum, n= 8, p<0.05 pre-specified threshold FDR, corrected for multiple comparisons using the Bonferroni correction; pFDR = 0.00625). This test complements the rGED analysis, as the 2D correlation calculation utilizes all connection pairs and does not require thresholding to create a binary network.

**Figure 10.**
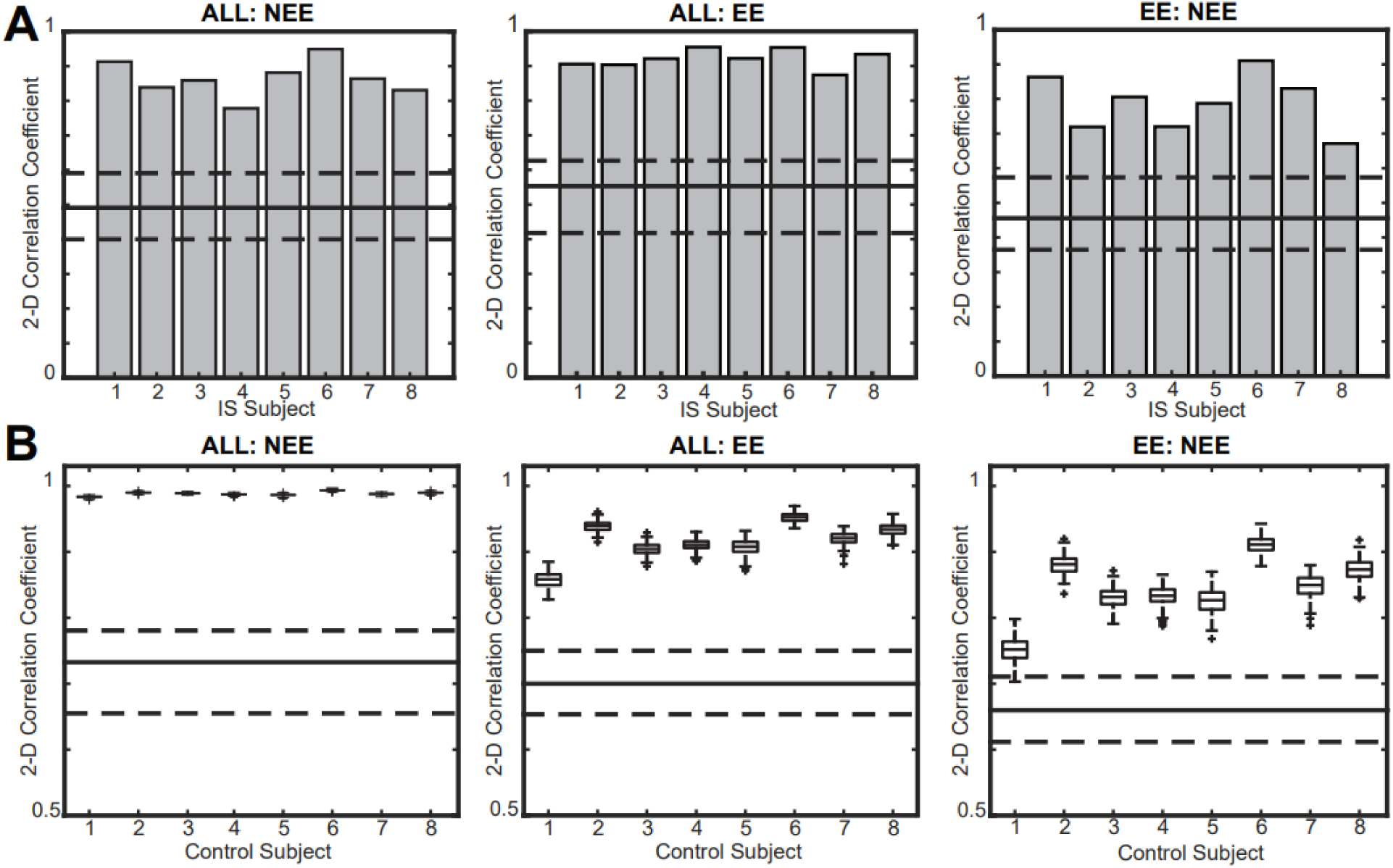
2D correlation coefficients for all intra-subject network comparisons are significantly higher than inter-subject comparisons in (A) IS subjects and (B) controls. Gray bars represent intra-subject comparisons for IS subjects, and box plots represent intra-subject values from 200 simulations in controls. The solid line represents the median of the inter-subject correlation values and the dashed lines represent the 25^th^ and 75^th^ percentiles.

### 3.6 Connectivity strength increases during IEDs in epilepsy subjects

Changes in connectivity strength in the presence and absence of an IED for each subject was done using three one-tailed Wilcoxon sign-rank tests. We tested the paired distributions of all 171 connectivity strengths for the following FCN comparisons within each subject: (1) EE > ALL, (2) ALL > NEE, and (3) EE>NEE. These comparisons were based on the simulated data in Figures 4, 5B, and 5D, which suggested that the EE network would have the highest connectivity strength, followed by the ALL network and then the NEE network. For all three network comparisons, this hypothesis held true for seven out of eight IS subjects (Table 2). In contrast, the sign-rank test yielded no significant differences in connectivity strength for any of the three network comparisons for the control subjects. The p-values for control subjects were equivalent across simulations with randomly placed IEDs, indicating that the specific timing of the IEDs did not affect the results.

**Table II.** P-values for statistical comparisons of connectivity strength between FCNs in individual subjects. The p-values from control subjects come from 200 simulations with spikes inserted at random times in the EEG record.

| *Cohort* | *Subject* | *FCN comparison* |  |  |
| --- | --- | --- | --- | --- |
|  |  | *EE>ALL* | *ALL>NEE* | *EE>NEE* |
| *IS* | 1 | **0** | 0.221 | **0** |
|  | 2 | **0** | **0.001** | **0** |
|  | 3 | 0.853 | **0.037** | 1 |
|  | 4 | **0** | **0.004** | **0** |
|  | 5 | **0** | **0.001** | **0.017** |
|  | 6 | **0** | **0** | **0.020** |
|  | 7 | **0** | **0.021** | **0.006** |
|  | 8 | **0** | **0** | **0** |
| *Controls* | 1 | 0.9590 ±0 | 0.9317 ±0 | 0.9951 ±0 |
| | 2 | 0.6156 ±0 | 0.0255 ±0 | 0.2771 ±0 |
| | 3 | 0.8938 ±0 | 0.9696 ±0 | 0.9782 ±0 |
| | 4 | 0.6366 ±0 | 0.3542 ±0 | 0.6889 ±0 |
| | 5 | 0.3525 ±0 | 0.9989 ±0 | 0.8491 ±0 |
| | 6 | 0.4402 ±0 | 0.2009 ±0 | 0.3651 ±0 |
| | 7 | 0.3827 ±0 | 0.6515 ±0 | 0.6954 ±0 |
| | 8 | 0.4520 ±0 | 0.2043 ±0 | 0.3104 ±0 |

## 4. Discussion

In this study, we investigated how the presence of IEDs impacts EEG-based FCNs. IS subjects have unique, patient-specific FCNs, while control subjects have more stereotyped network structures. In IS subjects, physiological IEDs do not significantly alter FCN structure, consistent with simulated IEDs in control subjects, which only alter FCN structure when the amplitude or spike burden are outside of normal physiological ranges. On the other hand, functional connectivity strength significantly increases during the occurrence of physiological IEDs in IS subjects, whereas the addition of simulated IEDs to normal EEG does not increase the connectivity strength. We conclude that these changes in connectivity strength in IS subjects are not spurious increases caused by the presence of the interictal spike waveform in the EEG. Overall, these findings suggest that an individual’s FCN structure remains stable during an IED, and that the changes in connectivity strength noted in epileptic subjects are likely due to the underlying pathophysiological networks of the disease, rather than simply the presence of the IED waveform itself. Furthermore, our analysis suggests that there is no need to mark and remove IEDs prior to calculating an FCN, as the functional connectivity measurements derived from EEG epochs containing IEDs do not differ significantly from those of interictal EEG epochs.

In control subjects with simulated IEDs, the amplitude of the IED affected FCN structure more than the spike burden did. Introducing simulated IEDs with high spike:background amplitude ratios caused major alterations to the FCN structure, indicated by a decrease in 2D correlation from 1.0 to nearly zero (Figure 5C). In comparison, high spike burdens of 100% led to a decrease in 2D correlation from 1.0 to 0.8, indicating that the increase in burden caused little change in the network structure (Figure 8).

Introducing simulated IEDs with physiological spike amplitudes and burdens derived from the EEG of IS subjects did not cause a substantial change in FCN structure. Across all focal IEDs detected in IS subjects, the average spike:background ratio was 2.62, while changes in the control subjects’ EE network structure emerged at a simulated spike:background ratio greater than 7. The consistency of network structure in IS subjects is shown in Figures 9 and 10, where the rGED and 2D correlations of intra-subject FCN comparisons are significantly higher than inter-subject comparisons. Increasing the spike burden also had minimal effect, resulting in a change of less than 5% in both 2D correlation and mean connectivity strength (Figure 7).

In addition to assessing the impact of IED amplitude and burden on FCNs, we investigated the effect of simulated IED location on the FCN of control subjects. Varying the location of focal IEDs produced similar effects as placing the focal spikes at F3, with minimal change noted in the FCN structure at low amplitudes. Increasing the simulated IED amplitudes past the physiological range resulted in increased connectivity strength and an altered FCN structure with long-range connections involving the IED focus, similar to Figure 4B. Overall, this result confirmed that IED location was not a crucial factor in our tests of IED amplitude and burden.

The shape of the IED waveform could also impact our results. Our simulated IED was modeled after the typical spike-wave complex associated with many types of epilepsy. However, there are many types of IED waveforms to consider, including isolated spikes (without a slow wave component), polyspikes with and without slow wave components, and paroxysmal fast activity (de Curtis et al. 2012). Here, we chose an IED waveform containing both a spike and a slow wave, as this had the greatest potential to affect the connectivity calculation. We also chose to include the slow wave in our simulation because the slow wave can drive changes in broadband EEG connectivity, and it has also been linked to hypsarrhythmia in IS subjects (Hrachovy and Frost 2003; Frost and Hrachovy 2005).

Prior to subject-specific FCN tests, we compared the FCN differences between the IS and control cohorts. Across all groupwise comparisons, we found that the median 2D correlation of control-control FCNs was approximately 0.75, while the median 2D correlation of control-IS and IS-IS comparisons was approximately 0.5 and 0.6, respectively (Figure 8). The significantly lower values for the control-IS comparison are in accordance with prior studies using EEG-based FCN, with reported differences in the global network characteristics of temporal lobe epilepsy and focal epilepsies compared to the healthy brain (Horstmann et al. 2010; Kramer and Cash 2012; Quraan et al. 2013). The low 2D correlation in IS-IS comparisons was likely due to the multifocal nature and the wide range of etiologies in IS, resulting in a unique network structure for each subject. Similar results were reported for EEG-based FCNs associated with absence seizures, where across patient correlations were lower than within patient comparisons (Taylor et al. 2013). We report a median 2D correlation between control subject FCNs of 0.75, higher than a previous study reporting cross-correlation values between healthy adult controls of approximately 0.5 (Chu-Shore et al. 2012). These differences in correlation could reflect a difference between the infant brain, as studied here, and the adult brain. Moreover, the correlation coefficient of 0.75 for control-control comparisons does not suggest an identical FCN structure across patients, but rather indicates that control subjects are more likely to exhibit stereotyped connectivity patterns than IS subjects.

Generally, the occurrence of IEDs in IS subjects was associated with a global increase in connection strength without changes in network structure. FCNs of epochs containing interictal spikes had the highest connectivity strength, followed by FCNs generated using all epochs, followed by FCNs lacking interictal spikes. These relationships are in agreement with previous studies, where the presence of interictal spikes resulted in increased connectivity strength in the medial temporal pole, hippocampus, amygdala, parahippocampal gyrus, olfactory gyrus, and gyrus rectus regions compared to baseline periods without spikes (Wilke et al. 2011; Coito et al. 2015). Of the eight IS subjects, seven had ALL, EE, and NEE FCN structures that were more similar to each other than they were to other subjects. For IS subject 8, the ALL and EE FCNs consisted of long-range connections emanating from the location of the IEDs, with higher connectivity strengths than all other epilepsy subjects. This is similar to the EE FCNs of controls with very high amplitude IEDs (Figure 4B). The rGED of the EE:NEE comparison for IS subject 8 was 0.24, compared to the other IS subjects who have rGED values of approximately 0.6 (Figure 9A). This low rGED value suggests a significant change in network structure during IEDs. Although IS subject 8 did not have a significantly larger spike:background amplitude ratio than the other IS subjects, they did have spike-wave discharges with significantly larger slow wave amplitudes than other IS subjects, which may explain this discordant finding. This could explain the similarity between this IS subject and the control subject simulations at high amplitudes, suggesting that the slow wave’s waveform is causing the changes in the measurement of FCN structure.

There are limitations to the methods presented here. We analyzed standard clinical EEG recordings with nineteen scalp electrodes to promote broad applicability to any clinical epilepsy center, as opposed to other studies that utilized high-density research EEG recordings (Coito et al. 2015, 2016; Mahmoudzadeh et al. 2016). Although our results were consistent with those from prior studies, the use of high-density electrode arrays would allow the use of additional graph theory metrics, whose accuracy depends on having a network with many nodes. In addition, our study is limited by a small sample size of sixteen subjects. Future studies can increase the cohort size for healthy controls, IS subjects, and non-IS epilepsy subjects to further test the validity and robustness of our findings. Other limitations of our study lie in the assumptions inherent to simulating IEDs. We used boundary element methods modelled on a template brain volume from the ICBM 152 atlas, which is based on young adults rather than infants. The properties of the simulated IED waveform were chosen to match that of a spike-wave complex, which does not reflect all IED types. Additionally, the model only simulated IEDs at a single location, which is simpler than the multifocal nature of IEDs in IS.

In future work, we plan to validate our findings by increasing the cohort size to include more subjects, including adults and older children, with both generalized and focal epilepsies. Due to the heterogeneous IEDs contained in IS subjects, we also plan to study different aspects of the spike-wave complex such as the effects of slow wave amplitude rather than the spike amplitude alone. Lastly, we plan to analyze the impact of different IED waveforms and the presence of multi-focal IEDs on functional networks. This work will elucidate the dynamic changes in functional connectivity and the robustness of FCNs in the presence of transient waveforms occurring over long spans of time.

## 5. Acknowledgements

The authors would like to thank the clinical epileptologists at CHOC Children’s for their contributions to this study. This work was supported in part by an Institute of Clinical and Translational Sciences UC Irvine-Children’s Hospital of Orange County Collaborative Grant and a Children’s Hospital of Orange County Pediatric Subspecialty Faculty Tithe Grant

## 9. Supplementary figures

**S1.**
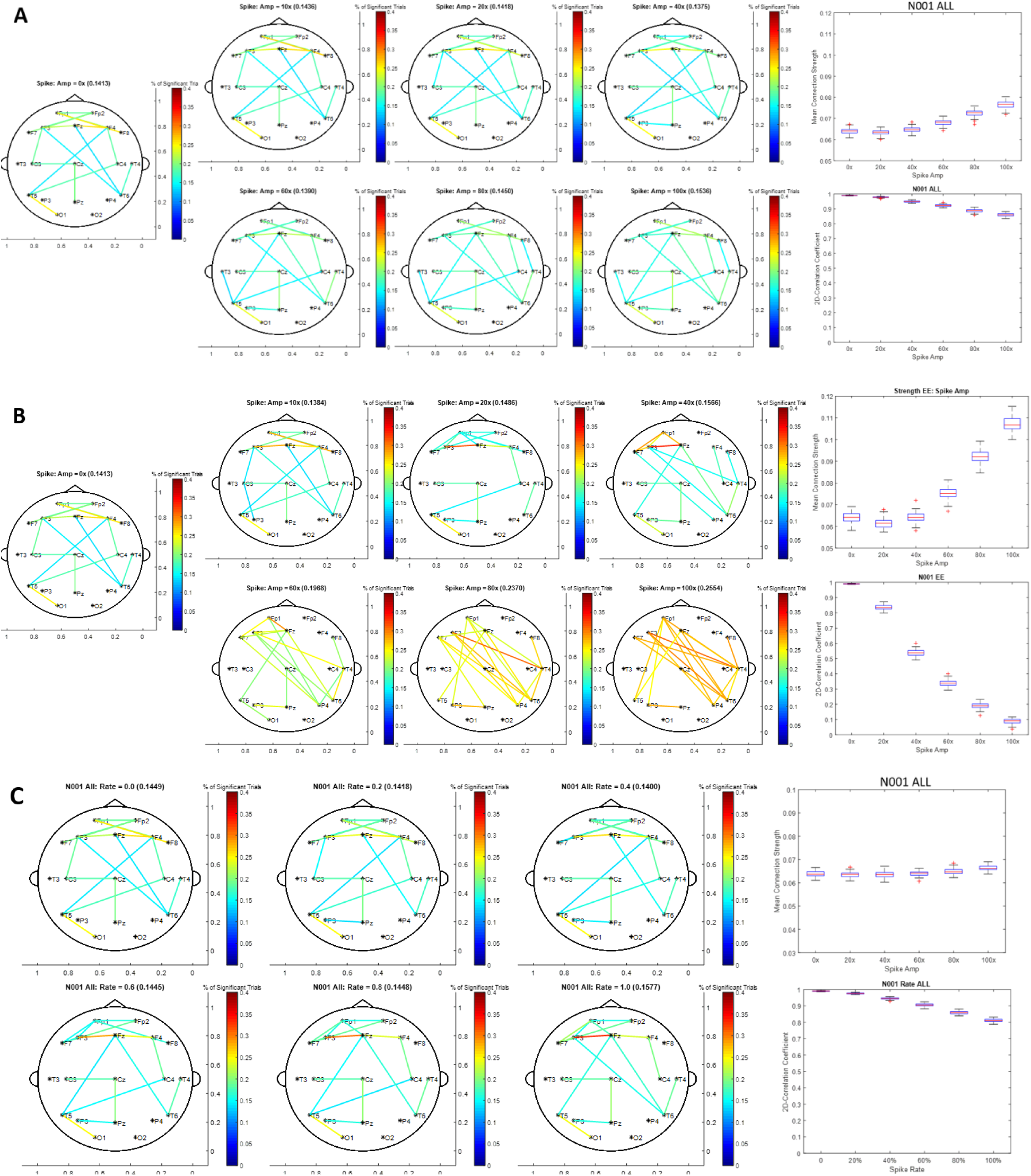
FCNs, connectivity strength tests, and network structure tests for control subject 1. Figures S1A and S1B are the ALL and EE network with simulated spikes with varying amplitudes at a fixed rate of 0.25. Figure S1C contains the ALL network with simulated spikes at varying rates at a fixed amplitude of 120 μV. FCNs and tests are done using the average of thirty different iterations of simulated IEDs.

**S2.**
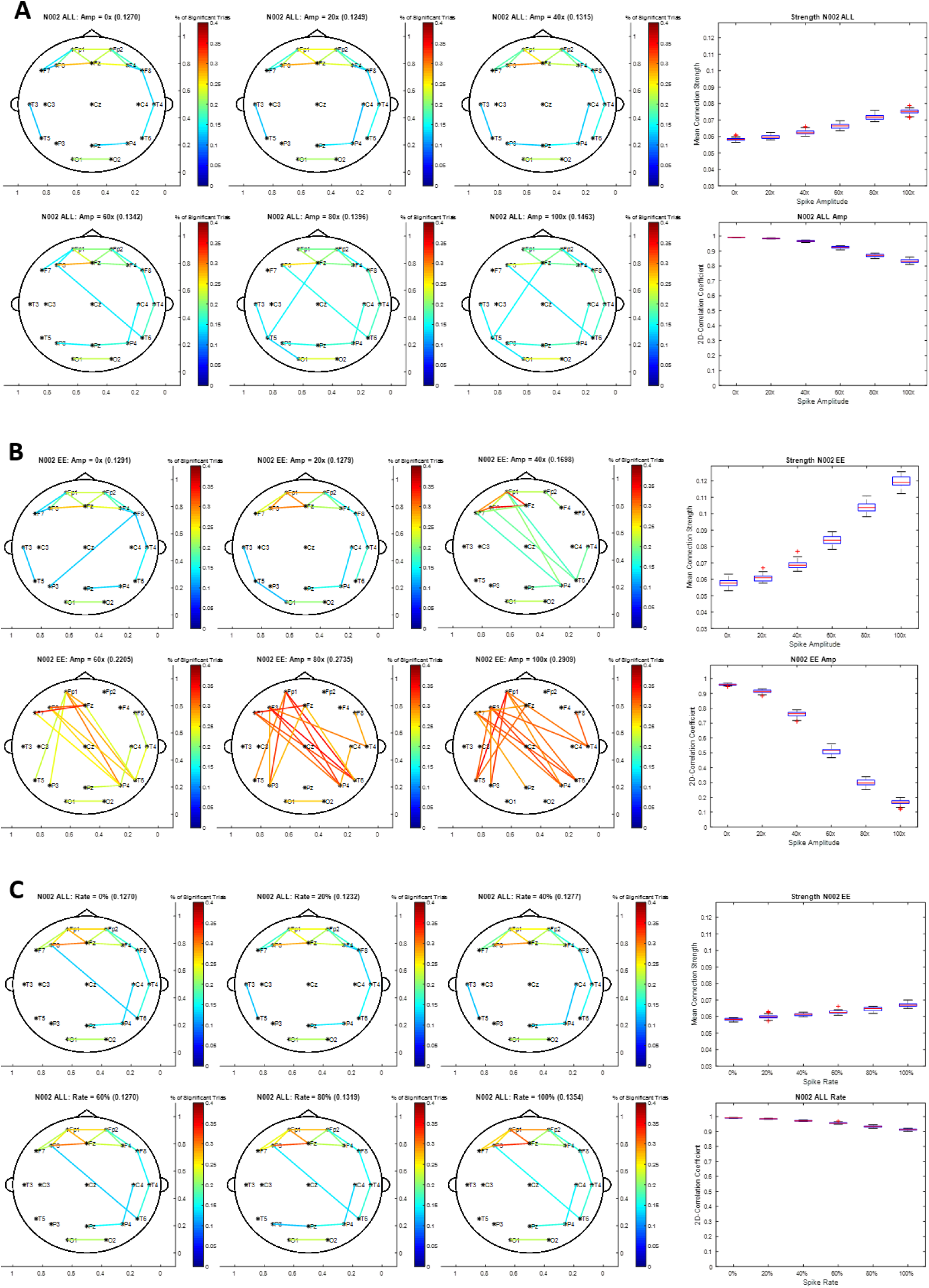
FCNs, connectivity strength tests, and network structure tests for control subject 2. Figures S2A and S2B are the ALL and EE network with simulated spikes with varying amplitudes at a fixed rate of 0.25. Figure S2C contains the ALL network with simulated spikes at varying rates at a fixed amplitude of 120 μV. FCNs and tests are done using the average of thirty different iterations of simulated IEDs.

**S3.**
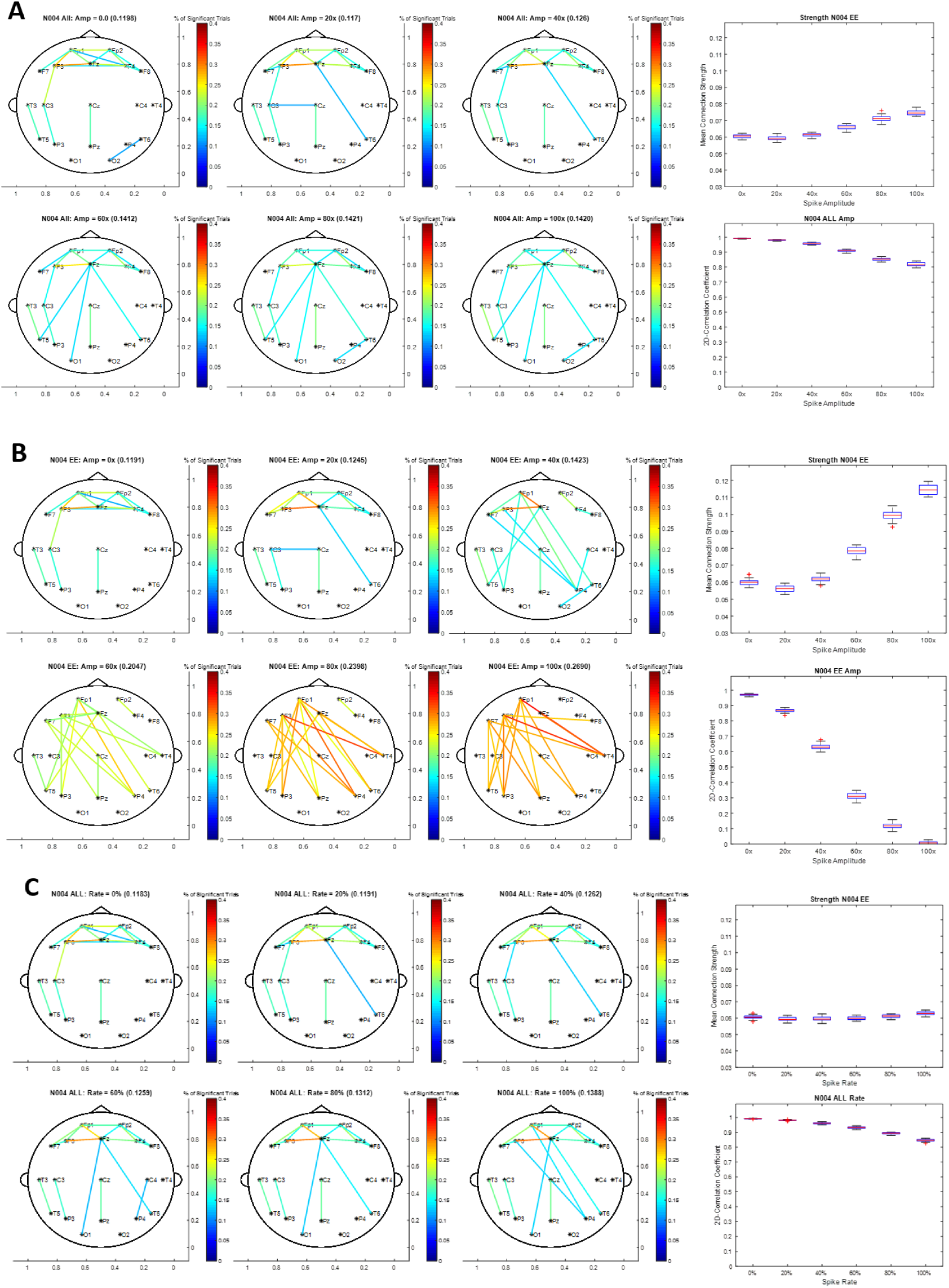
FCNs, connectivity strength tests, and network structure tests for control subject 3. Figures S3A and S3B are the ALL and EE network with simulated spikes with varying amplitudes at a fixed rate of 0.25. Figure S3C contains the ALL network with simulated spikes at varying rates at a fixed amplitude of 120 μV. FCNs and tests are done using the average of thirty different iterations of simulated IEDs.

**S4.**
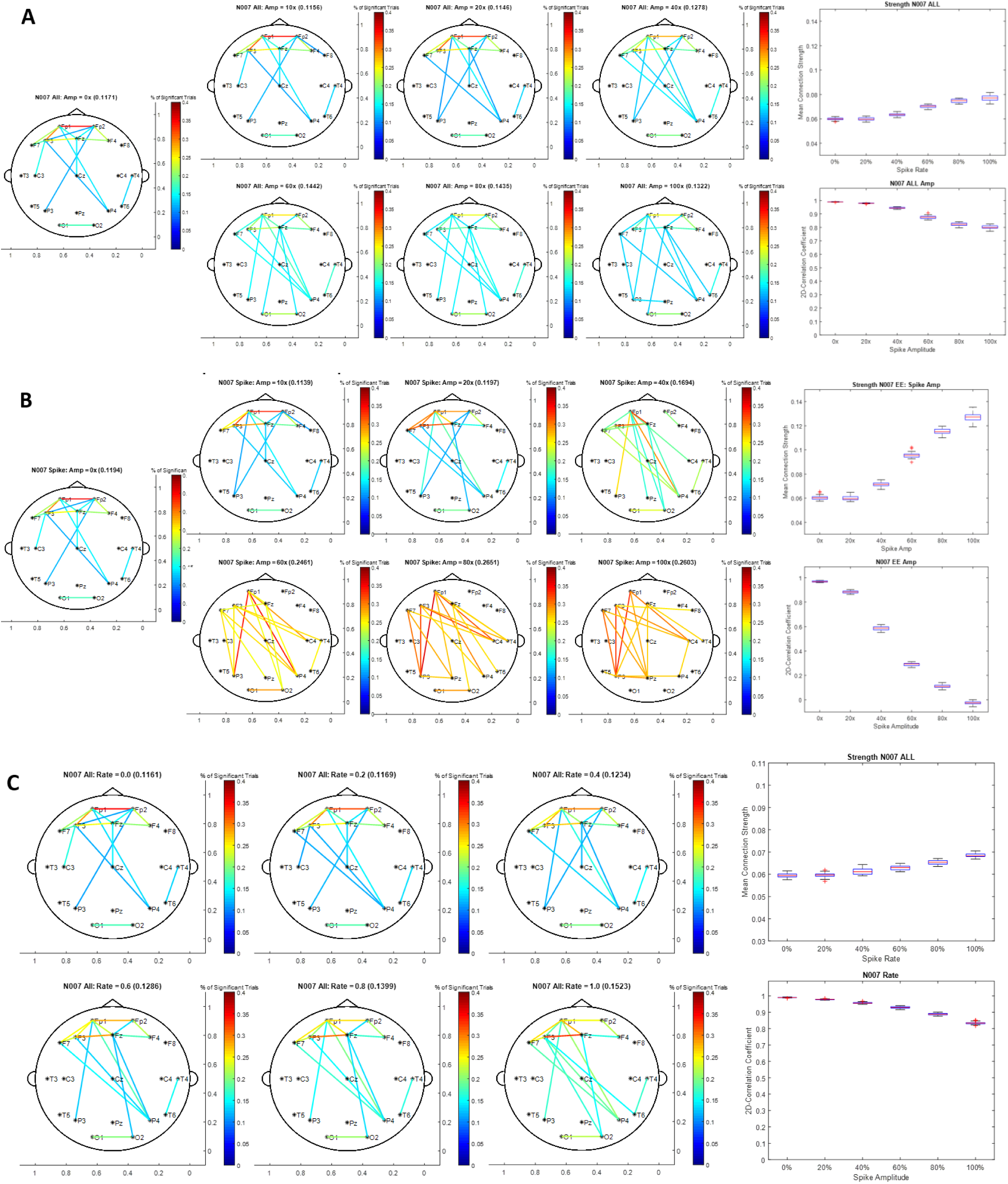
FCNs, connectivity strength tests, and network structure tests for control subject 4. Figures S4A and S4B are the ALL and EE network with simulated spikes with varying amplitudes at a fixed rate of 0.25. Figure S4C contains the ALL network with simulated spikes at varying rates at a fixed amplitude of 120 μV. FCNs and tests are done using the average of thirty different iterations of simulated IEDs.

**S5.**
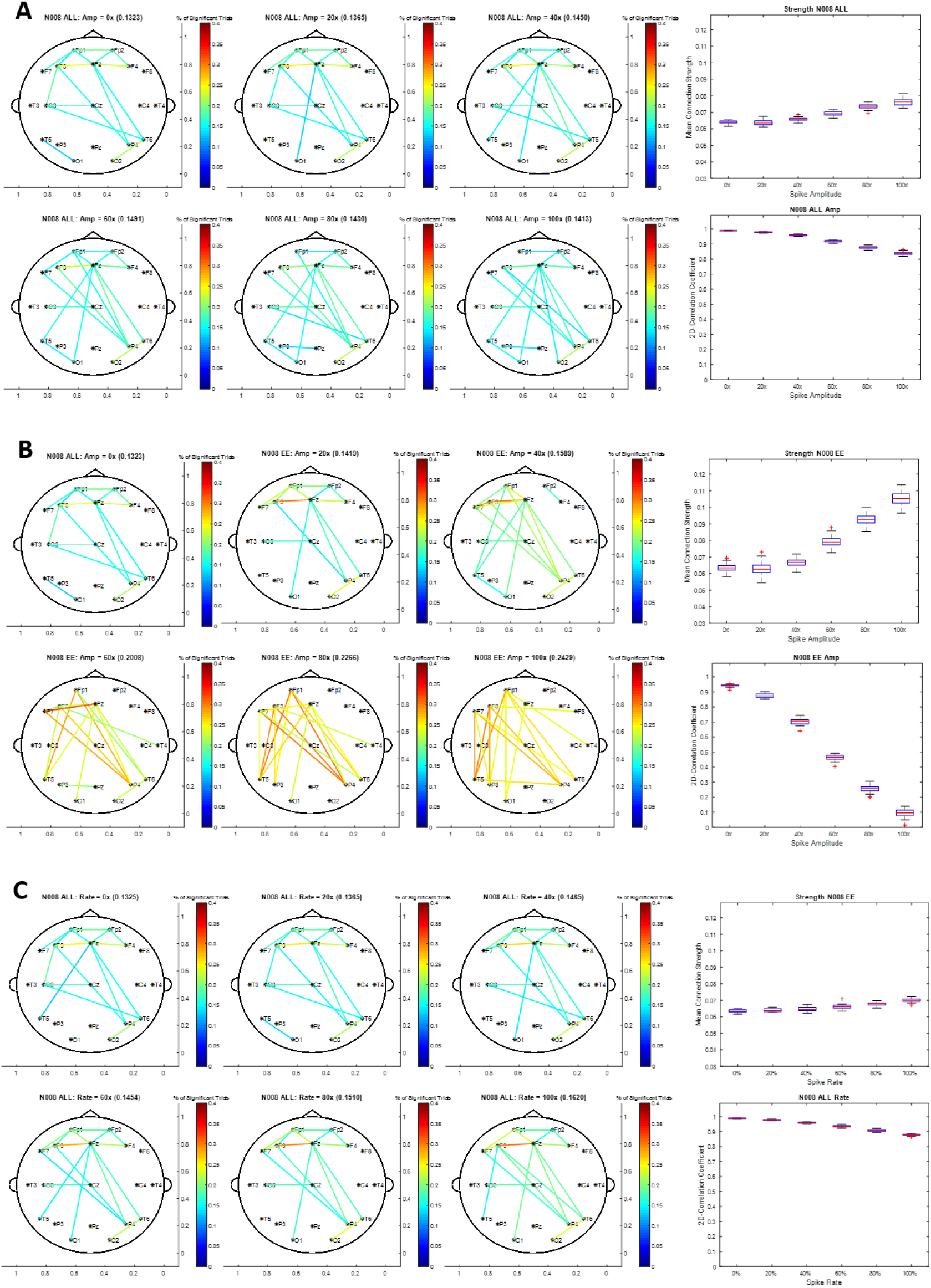
FCNs, connectivity strength tests, and network structure tests for control subject 5. Figures S5A and S5B are the ALL and EE network with simulated spikes with varying amplitudes at a fixed rate of 0.25. Figure S5C contains the ALL network with simulated spikes at varying rates at a fixed amplitude of 120 μV. FCNs and tests are done using the average of thirty different iterations of simulated IEDs.

**S6.**
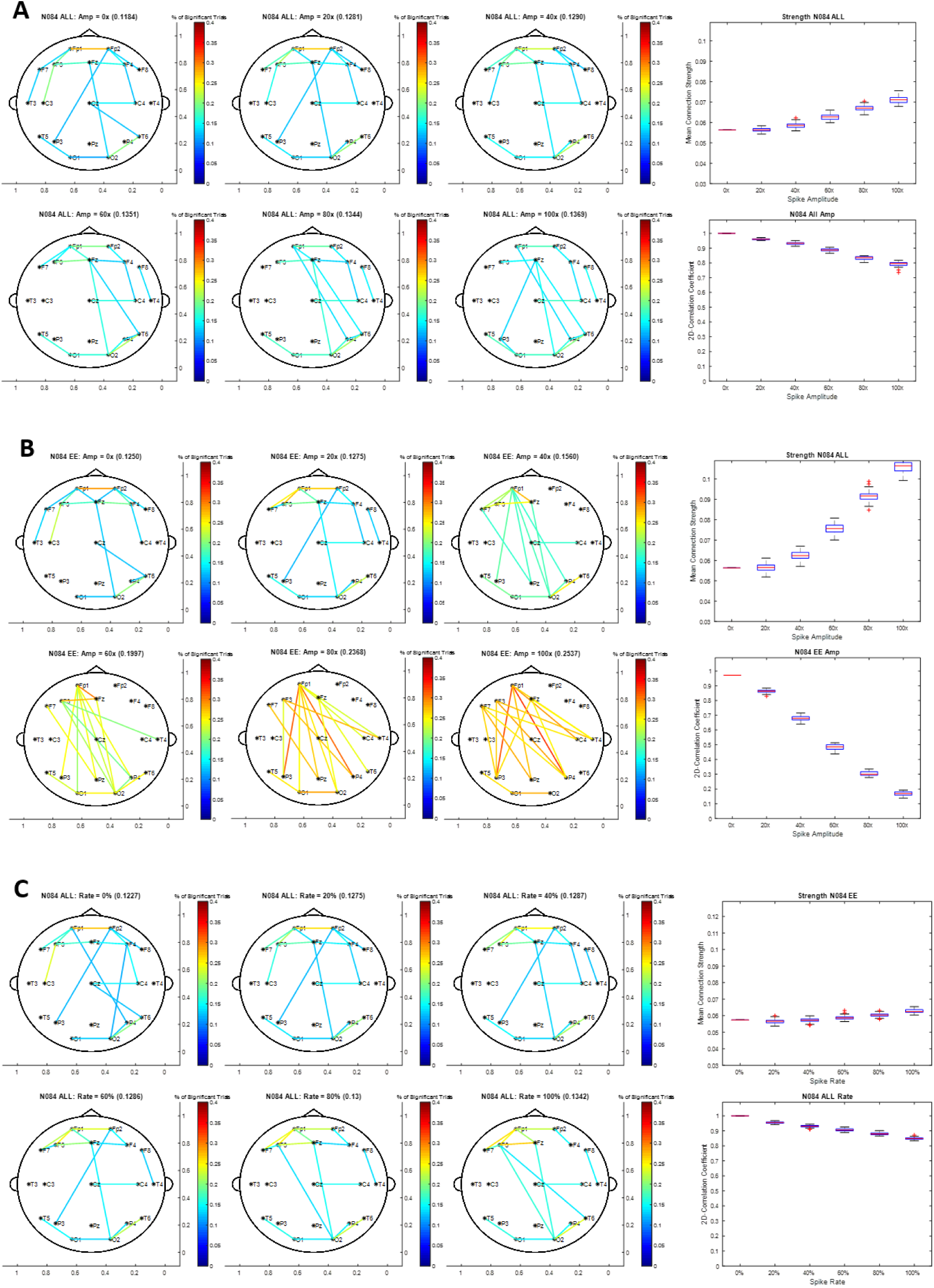
FCNs, connectivity strength tests, and network structure tests for control subject 7. Figures S7A and S7B are the ALL and EE network with simulated spikes with varying amplitudes at a fixed rate of 0.25. Figure S7C contains the ALL network with simulated spikes at varying rates at a fixed amplitude of 120 μV. FCNs and tests are done using the average of thirty different iterations of simulated IEDs.

**S7.**
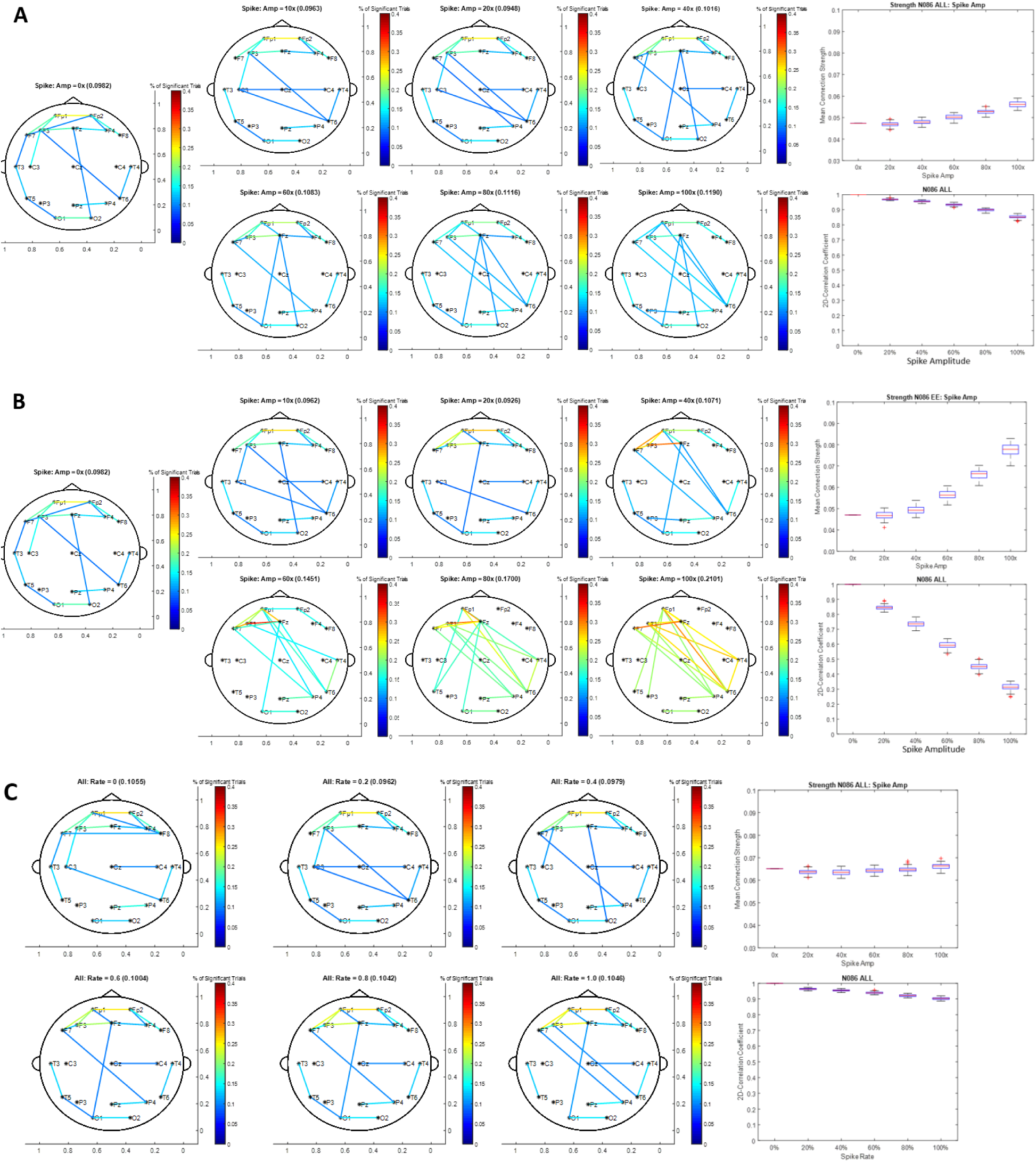
FCNs, connectivity strength tests, and network structure tests for control subject 8. Figures S8A and S8B are the ALL and EE network with simulated spikes with varying amplitudes at a fixed rate of 0.25. Figure S8C contains the ALL network with simulated spikes at varying rates at a fixed amplitude of 120 μV. FCNs and tests are done using the average of thirty different iterations of simulated IEDs.

